# Genetic diversity and population structure of the endangered endemic species *Paeonia decomposita* from China and implications for its conservation

**DOI:** 10.1101/860734

**Authors:** Shi-Quan Wang

## Abstract

*Paeonia decomposita*, endemic to China, has important ornamental, medicinal and economic value and is regarded as a threatened endangered plant. The genetic diversity and structure have seldom been described. A conservation management plan is not currently available. In present study, 16 pairs of SSR primers were used to evaluate genetic diversity and population structure. A total of 122 alleles were obtained with a mean of 7.625 alleles per locus. The expected heterozygosity (He) varied from 0.043 to 0.901 (mean 0.492). Moderate genetic diversity (He=0.405) among populations were revealed, with Danba identified as the center of genetic diversity. Mantel tests revealed a significant positive correlation between geographic and genetic distance among populations (r=0.592, P=0.0001), demonstrating consistency with the isolation by distance model. Analysis of molecular variance (AMOVA) results indicated that the principal genetic variation existed within populations (73.48%) rather than among populations (26.52%). Bayesian structure analysis and principal coordinate analysis (PCoA) supported classification of the populations into three clusters. Based on the level of observed genetic diversity, three management unints were proposed as conservation measures. The results will be beneficial for the conservation and exploitation of the species, providing a theoretical basis for further research on its evolution and phylogeography.

**Hightlights:**

1. Genetic diversity among populations was moderate in *Paeonia decomposita*
2. There is significant positive correlation between geographic and genetic distance among populations, consistent with the isolation by distance model
3. Principal genetic variation existed within populations rather than among populations.
4. The populations divided into three clusters.
5. Three management unints were proposed as conservation measures.

## 1. Introduction

The genus *Paeonia* L. (Paeoniaceae) includes 32 woody and herbaceous species, mainly distributed in the northern hemisphere. *Paeonia* is divided into three sections: Sections *Onaepia*, *Moutan* and *Paeonia* (Stern, 1946; Hong, 2010). The *Moutan* section comprises nine wild species native and endemic to China (Hong, 2010)and generally termed Mudan or tree peonies in Chinese. In China, Mudan is regarded as the ‘King of Flowers’ and plays an important role in pharmaceutical exploitation of the plant in addition to having ornamental value (Stern, 1946; Cheng, 2007). Seed oil can be extracted from peony seeds, which contain fatty acids and so they have become important woody oil crops (Zhang *et al*., 2015).

*Paeonia decomposita* Handel-Mazzetti is a species from the *Moutan* section, which principally occurs in remote mountain areas in northwest Sichuan Province, an indigenous and endemic species to China, with a sporadic and narrow distribution and small population size. It is cross-pollinated by insects (Yang *et al*., 2015) and propagated by seeds (Cheng *et al*., 1997). In the past, *P. decomposita* has consisted of two subspecies: *P. decomposita* subsp. *rotundiloba* and *P. decomposita* subsp. *decomposita* (Hong 1997; Hong and Pan, 1999; Hong *et al*., 2001). Based on morphological traits and molecular data, they are considered independent species (Hong, 2010; Hong, 2011).

*P. decomposita* is a famous ornamental flower on account of its large, showy, colorful and fragrant flowers. Thus, local people collect plants to use in ornamental gardening. It is also a traditional medicinal plant because its root bark (‘Dan Pi’ in Chinese) is used as a traditional Chinese medicine, having many therapeutic properties, for example, clearing heat, cooling blood, activating blood flow and removing blood stasis (Li *et al*., 2012). It has recently become an important woody oilseed plant. The mean kernel oil content was found to be 32.23±1.96%, consisting of seven fatty acids. Between 91.94 and 93.70% of the oil was found to be unsaturated fatty acids, with linolenic acid accounting for 40.45∼47.68% (Yang *et al*., 2015). The extracted oil from seeds can be utilized as oleochemicals, cosmetics and medicines (Han *et al*., 2014). Therefore, *P. decomposita* is considered to be not only an ornamental plant, but also an important officinal plant and a valuable woody oil crop.

Due to multiple threats including habitat damage, excessive harvesting of seeds, misuse of root-bark in traditional Chinese medicine and naturally poor regeneration, its natural habitats have become increasingly fragmented, the natural size of the population and individual numbers having decreased dramatically, resulting in a significant loss of genetic resource. Currently, most populations are small, fragmented and scattered, increasing the probability of inbreeding and the potential for genetic drift. Also, low seed production, difficult seedling renewal and the lack of a specific mechanism for long-distance seed dispersal has resulted in poor population regeneration because many communities are short of seedlings and saplings. In accordance with their distribution, biological characteristics and survival status, *P. decomposita* has been listed as a rare, endangered and threatened plant on the brink of extinction (Hong *et al*., 2017). Its conservation is therefore urgently needed.

Genetic resource conservation and plant breeding programs require an evaluation of the genetic diversity and structure of the endangered species (Cohen *et al*., 1991; Ouborg, 2010). It is difficult to conduct conservation strategies for this plant due to a lack of genetic background knowledge.

It is crucial that appropriate molecular markers are used to precisely estimate genetic diversity in order to protect valuable wild tree peonies and better understand and sustainably utilize them as a genetic resource. In the past, researchers have used various molecular markers to study the genetic relationships among the species in Section *Moutan* (Zhou *et al*., 2003; Lin *et al*., 2004; Meng *et al*., 2004; Zhao *et al*., 2004; Zhao *et al*., 2008). Compared with AFLP, SRAP and RAPD, SSR markers have the significant advantages of co-dominance, wide distribution, high transferability, high polymorphism, high reproducibility, high reliability combined with relatively low expense (Varshney *et al*. 2005; Agarwal *et al*. 2008), generally being regarded as ideal molecular markers. They have been widely employed to study genetic diversity, population structure and the genetic relationship of different plant species (Kumar *et al*., 2014; Wu *et al*., 2016; Aboukhalid *et al*., 2017; Litkowiec *et al*., 2018; Ni *et al*., 2018), including tree peonies (Yuan *et al*., 2012; Yu *et al*., 2013; Xu *et al*., 2016).

To date, the study of *P. decomposita* has been limited to the genetic relationships among species and the genetic diversity of inter simple sequence repeats (ISSRs) (Tong *et al*., 2016), with no studies exploring the genetic diversity of SSRs, or the genetic relationship or population structure of this important woody oilseed species. No breeding plan has been established from which to select an optimum germplasm or resource conservation strategy, preventing conservation of *P. decomposita*. Thus, an accurate understanding of the population structure and genetic diversity of *P. decomposita* is urgently required.

Accordingly, given its great value within medicine, industry and in ornamental applications, a genetic study of the plant was conducted. In the present study, we first selected 16 pairs of polymorphic SSR markers from tree peonies used throughout history. From these, the genetic variation among and within populations was evaluated, the genetic diversity analyzed and the population structure among and within populations estimated, providing crucial information for establishing an appropriate conservation and management strategy of genetic germplasm resource and deployment of these resources with plans for a future directive breeding strategy.

## 2. Materials and Methods

### 2.1 Plant materials

A total of 258 individual plants was sampled from eleven natural populations of *P. decomposita* across almost the complete regional distribution of China in 2017 prior to the flowering season. Ten–40 individual plants that were at least 10 m apart were sampled from each population. Details of the sampling are listed in Table 1. Fresh, tender and healthy leaves were individually sampled in the wild, then immediately placed in plastic, sealed bags and dried using colored silica gel then stored at −20°C until DNA was isolated.

**Table 1.** The sampling information of 11 populations of *Paeonia decomposita*

| Population | Population ID | Geographical coordinate |  | Altitude (m) | Sample size |
| --- | --- | --- | --- | --- | --- |
|  |  | Latitude(°N) | Longitude(°E) |  |  |
| Danba 1 | DB1 | 30.96503486°N | 101.8592443°E | 2819 | 21 |
| Danba 2 | DB2 | 30.84260025°N | 101.9128609°E | 2490 | 28 |
| Jinchuan 1 | JC1 | 31.48171608°N | 102.0683160°E | 2212 | 13 |
| Jinchuan 2 | JC2 | 31.31824367°N | 102.0190106°E | 2135 | 24 |
| Jinchuan 3 | JC3 | 31.28406350°N | 102.0008489°E | 2865 | 20 |
| Jinchuan 4 | JC4 | 31.36665014°N | 101.9824393°E | 2240 | 22 |
| Jinchuan 5 | JC5 | 31.46720113°N | 102.1038017°E | 2332 | 18 |
| Maerkang 1 | M1 | 31.99887580°N | 102.0183703°E | 2498 | 40 |
| Maerkang 2 | M2 | 31.88084028°N | 102.2571444°E | 2690 | 40 |
| Maerkang 3 | M3 | 31.92747578°N | 102.1097246°E | 2566 | 22 |
| Maerkang 4 | M4 | 31.91673120°N | 102.1856441°E | 2647 | 10 |

### 2.2 DNA extraction and PCR amplification

Total genomic DNA from each accession was extracted from the leaves using a Plant DNeasy Mini kit (Tiangen Biotech, Beijing, China) in accordance with the manufacturer’s instructions. DNA concentration and quality were measured using spectrophotometry and gel electrophoresis on 1% agarose, respectively. Extracted DNA was diluted to a working concentration of 50 ng/µl then stored at –20°C until required. Primers previously documented and developed for tree peonies were selected for screening. These primers were screened on representative samples and after initial screening, 16 polymorphic microsatellite primer pairs (Wang *et al*., 2009; Homolka *et al*., 2010; Hou *et al*., 2011; Hou *et al*., 2011; Zhang *et al*., 2012; Gao *et al*., 2013; Gilmore *et al*., 2013; Wang *et al*., 2013; Cai 2015) (Table 2) producing a high degree of polymorphism and high level of amplification were selected for subsequent analysis.

**Table 2.**
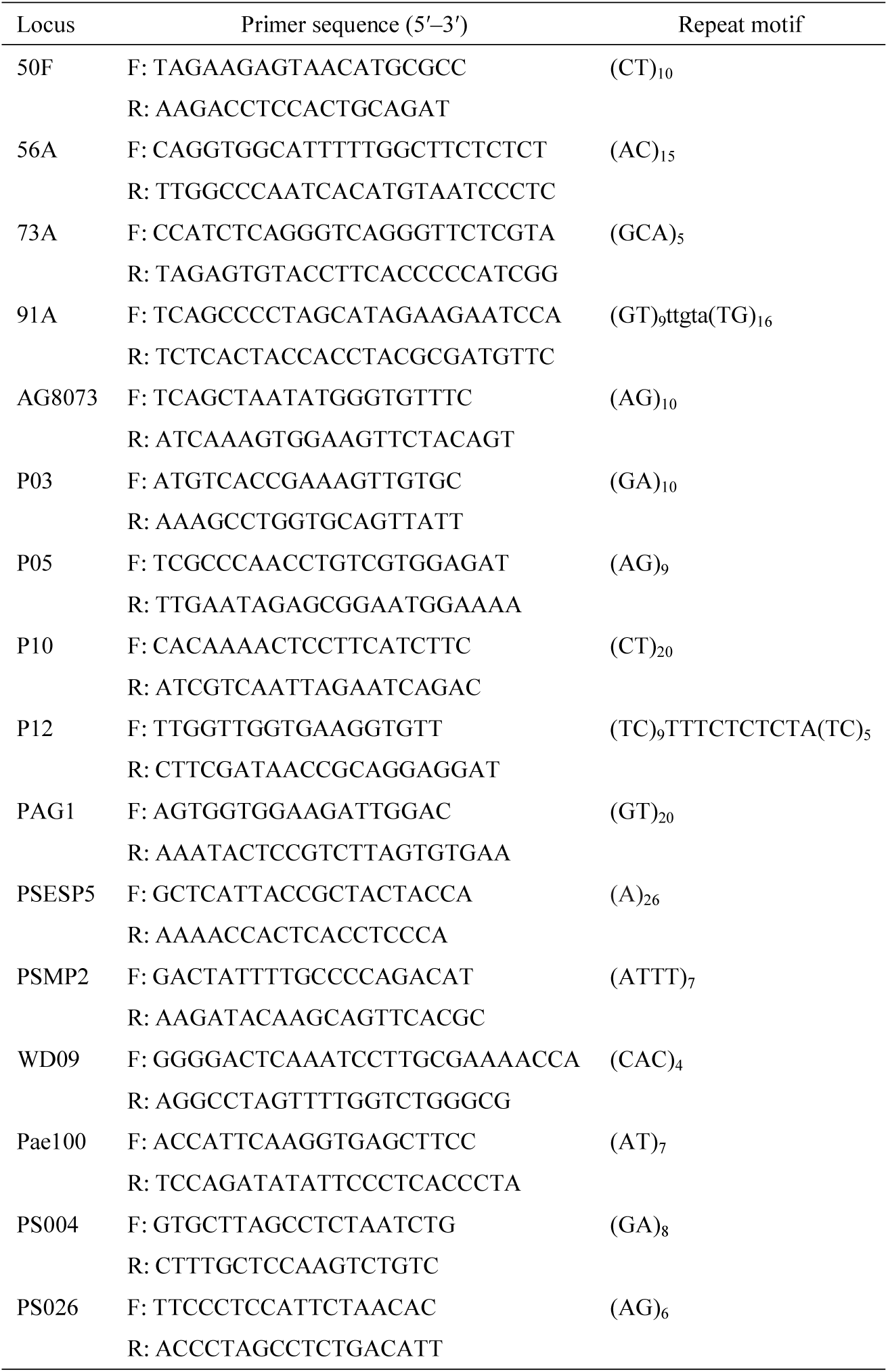
Characteristics of 16 polymorphic microsatellite primers

SSR-PCR amplification reactions were conducted using a total volume of 10 µl consisting of 5µl of 2×Taq PCR MasterMix (Tiangen, Beijing, China) (0.1 U/µl Taq DNA Polymerase, 0.5 mM each of dNTPs, 20 mM Tris-HCl, 100 mM KCl, 3 mM MgCl_2_), 1 µl of genomic DNA template (50 ng/µl) from each accession, 3 µl of ddH_2_O, and 1 μl of each primer. PCR amplification was conducted using a Bio-Rad thermal cycler (Applied Biosystems) with either of 2 different cycling protocols, as follows: 1. Pre-denaturation at 95°C for 5 min, followed by 10 cycles of denaturation at 95°C for 30 s, annealing at 62-52°C for 30 s (1°C drop for each cycle) and extension at 72°C for 30 s. 2. 25 cycles of denaturation at 95°C for 30 s, annealing at 52°C for 30 s then extension at 72°C for 30 s, followed by a final extension at 72°C for 20 min. All PCR products were genotyped using capillary electrophoresis on an ABI 3730XL DNA Analyzer. The alleles of all loci were scored relative to LIZ 500, an internal product size standard, with the aid of GeneMarker Version 4.0 (Softgenetics, USA).

### 2.3 Data analyses

The following genetic polymorphism parameters of the 11 populations were computed using GenAlEx 6.5 software (Peakall and Smouse, 2012): the number of polymorphic loci (NPL), percentage of polymorphic loci (PPL), the number of observed alleles per locus (N_a_), the number of effective alleles (N_e_), Shannon’s Information Index (I), observed heterozygosity (H_o_), expected heterozygosity (H_e_), fixation index (F), the inbreeding coefficient (F_is_) across populations and across loci, genetic differentiation coefficient (F_st_), gene flow (N_m_), pairwise Nei’s genetic distance (NGD), genetic identity (NGI) between populations and F-statistics (F_is_, F_it_ and F_st_) for each individual microsatellite locus across all populations. Polymorphic information content (PIC) was computed using Cervus 3.0 software (Kalinowski, *et al*., 2007), with allelic richness (A_r_) and intra-population inbreeding coefficients (F_is_) calculated using FSTAT 2.9.3.2 software (Goudet, 2001). Geographic distances (GGD) between populations were calculated using the online website http://www.hhlink.com/%e7%bb%8f%e7%ba%ac%e5%ba%a6 based on the geographical coordinates of the collection sites of each population. Hardy Weinberg equilibrium (HWE) deviation between microsatellites was tested using GenAlEx 6.5. A Mantel test was conducted to check correlations between matrices of genetic distances (GD) and geographic distances using GenAlEx 6.5. Population genetic structure was analyzed using a Bayesian clustering analysis method conducted in Structure 2.3.4 software (Pritchard *et al*., 2000). A total of ten independent runs (K=2–10) were performed with a run length of 1 × 10^5^ Markov Chain Monte Carlo (MCMC) replicates after a burn-in period of 1 × 10^5^ iterations in an admixture model with correlated allele frequency. The ΔK method (Evanno *et al*., 2005) was employed to select the most appropriate K value and optimal number of genetic clusters on Structure Harvester V6.0 software (Earl and vonHoldt, 2012). Principle coordinate analysis (PCoA) was used with GenAlEx 6.5 to evaluate the genetic relationships between populations in the light of Nei’s genetic distance matrices between all pairwise populations.

Analysis of molecular variance (AMOVA) was further used to evaluate the hierarchical distribution of genetic variation among regions, within regions, among populations, within populations, and among populations within regions using GenAlEx 6.5. An unweighted pair group method with arithmetic mean (UPGMA) dendrogram was generated from cluster analysis with 1000 bootstrap replications using PHYLIP v3.67 software (Felsenstein, 2007) on the basis of Nei’s genetic distance, which was then used to assess the genetic relationships among populations.

## 3. Results

### 3.1 SSR marker polymorphism

In this study, representative samples from 11 populations of *P. decomposita* were randomly chosen to perform primer prescreening for genetic polymorphism. Sixteen pairs of primers which were successfully amplified and producing clear, unambiguous, reproducible banding patterns were further used for PCR amplification, polymorphism identification and genetic diversity analysis of 258 samples (Table 3). A total of 122 alleles at 16 polymorphic microsatellite loci were amplified across 258 individual plants of 11 natural populations, the number of observed alleles per locus (Na) varying greatly among loci, from 2 alleles (locus PSMP2) to 20 alleles (locus PAG1) (mean of 7.625). The number of effective alleles per locus (Ne) ranged from 1.045 (for primer PSMP2) to 9.929 (for primer PAG1) (mean of 3.208). Observed heterozygosity per locus (Ho) ranged from 0.027 (locus WD09) to 0.992 (locus 73A) (mean of 0.385), whereas expected heterozygosity (He) ranged from 0.043 (locus PSMP2) to 0.901(locus PAG1), a mean of 0.492. The polymorphism information content (PIC value) of primers varied from 0.042 (locus PSMP2) to 0.891 (locus PAG1) with a mean of 0.456.

**Table 3.** Performance of the microsatellite markers on 258 samples across 11 populations of *Paeonia decomposita* in China.

| Locus | $N_a$ | $N_e$ | $I$ | $H_o$ | $H_e$ | $F$ | $PIC$ | $A_r$ | $F_{is}$ | $F_{st}$ | $N_m$ |
| --- | --- | --- | --- | --- | --- | --- | --- | --- | --- | --- | --- |
| 50F | 3 | 1.293 | 0.440 | 0.125 | 0.227 | -0.045 | 0.209 | 3 | -0.026 | 0.430 | 0.331 |
| 56A | 15 | 5.355 | 1.926 | 0.395 | 0.815 | 0.389 | 0.789 | 14.552 | 0.374 | 0.195 | 1.033 |
| 73A | 3 | 2.548 | 1.003 | 0.992 | 0.609 | -0.798 | 0.53 | 3 | -0.784 | 0.091 | 2.502 |
| 91A | 13 | 7.624 | 2.214 | 0.553 | 0.871 | 0.310 | 0.856 | 12.981 | 0.308 | 0.138 | 1.556 |
| AG8073 | 3 | 1.252 | 0.408 | 0.113 | 0.202 | 0.129 | 0.189 | 3 | 0.148 | 0.302 | 0.577 |
| P03 | 3 | 1.803 | 0.690 | 0.401 | 0.446 | -0.091 | 0.359 | 3 | -0.182 | 0.255 | 0.732 |
| P05 | 5 | 2.145 | 0.851 | 0.416 | 0.535 | 0.128 | 0.425 | 4.848 | 0.081 | 0.175 | 1.175 |
| P10 | 7 | 2.443 | 1.183 | 0.486 | 0.592 | 0.003 | 0.547 | 6.883 | -0.001 | 0.146 | 1.467 |
| P12 | 15 | 5.388 | 1.977 | 0.711 | 0.816 | -0.038 | 0.793 | 14.745 | -0.035 | 0.139 | 1.552 |
| PAG1 | 20 | 9.929 | 2.540 | 0.590 | 0.901 | 0.155 | 0.891 | 19.972 | 0.154 | 0.153 | 1.385 |
| PSESP5 | 4 | 1.122 | 0.272 | 0.093 | 0.109 | 0.033 | 0.106 | 4 | 0.062 | 0.084 | 2.735 |
| PSMP2 | 2 | 1.045 | 0.106 | 0.036 | 0.043 | -0.019 | 0.042 | 2 | -0.002 | 0.151 | 1.400 |
| WD09 | 3 | 1.049 | 0.128 | 0.027 | 0.046 | 0.245 | 0.046 | 3 | 0.245 | 0.211 | 0.936 |
| Pae100 | 8 | 1.484 | 0.762 | 0.187 | 0.327 | 0.365 | 0.313 | 7.886 | 0.284 | 0.235 | 0.815 |
| PS004 | 12 | 4.682 | 1.848 | 0.687 | 0.788 | -0.040 | 0.762 | 11.865 | -0.031 | 0.139 | 1.554 |
| PS026 | 6 | 2.161 | 0.868 | 0.345 | 0.538 | 0.027 | 0.431 | 5.689 | 0.020 | 0.250 | 0.751 |
| mean | 7.625 | 3.208 | 1.076 | 0.385 | 0.492 | 0.047 | 0.456 | 7.526 | 0.038 | 0.193 | 1.281 |

$N_a$ : The observed number of allele.
$N_e$ : The effective number of alleles.
$I$ : Shannon's information index.
$H_o$ : Observed heterozygosity.
$H_e$ : Expected heterozygosity.
$F$ : Fixation index.
$PIC$ : Polymorphism information content.
$A_r$ : Allelic richness.
$F_{is}$ : Inbreeding coefficient among individuals within populations.

### 3.2 Population genetic diversity

At the population level, genetic diversity indices (in terms of NPL, PPL, Na, Ne, I, Ho, He, F) varied across populations of *P. decomposita*, as listed in Table 4. The number of polymorphic loci (NPL) in each population (pop.) varied from 9 (pops. JC5, M1, M4) to 14 (pops. DB1, M3). On average, the percentage of polymorphic loci (PPL) across eleven populations was high (78%) and ranged from 64% for JC5 to 93% for DB1, the majority of populations (9/11) slightly over 70%. The number of observed alleles (Na) per population varied from 2.563 (pop. M4) to 4.813 (pop. DB2), with a mean of 3.637. The number of effective alleles (Ne) across all populations was 2.322, varying from 1.811 (pop. M1) to 2.813 (pop. DB2). Mean heterozygosity (He) and observed heterozygosity (Ho) across all populations was in the range 0.329 (pop. M1) to 0.538 (pop. DB1) and 0.314 (pop. M2) to 0.464 (pop. DB2), with a mean of 0.405 and 0.394, respectively. The mean value of Shannon’s Information Index (I) was 0.777 over a range of 0.58 (pop. M1) to 1.017 (pop. DB1). The fixation index (F) averaged 0.032, ranging from -0.160 (pop. JC5) to 0.154 (pop. DB1) at locus and population levels. Most of the loci accord with Hardy–Weinberg Equilibrium (HWE), but some populations partly were not in HWE, especially the populations DB2 and M3 which indicated a lot of loci deviating from HWE (7 and 8 loci respectively), showing a pan fusion population structure. Loci 56A and 73A deviated from HWE in all populations.

**Table 4.** Genetic variation of the 11 populations in *Paeonia decomposita*

| Pop | $N_a$ | $N_e$ | $I$ | $H_o$ | $H_e$ | $F$ | $PIC$ | $A_r$ | $NPL$ | $PPL$ | $F_{is}$ | $HWE$ |
| --- | --- | --- | --- | --- | --- | --- | --- | --- | --- | --- | --- | --- |
| DB1 | 4.313 | 2.714 | 1.017 | 0.456 | 0.538 | 0.154 | 0.456 | 3.731 | 14 | 93% | 0.178 | 56A<sup>*</sup>, 73A<sup>***</sup>, 91A<sup>***</sup>, PAG1<sup>*</sup>, PSESP5<sup>***</sup>, PS004<sup>**</sup> |
| DB2 | 4.813 | 2.813 | 0.989 | 0.464 | 0.486 | 0.053 | 0.464 | 3.892 | 12 | 86% | 0.066 | 56A<sup>**</sup>, 73A<sup>***</sup>, 91A<sup>*</sup>, AG8073<sup>*</sup>, P05<sup>***</sup>, PAG1<sup>***</sup>, Pae100<sup>*</sup> |
| JC1 | 3.875 | 2.616 | 0.826 | 0.437 | 0.407 | -0.045 | 0.437 | 3.666 | 12 | 75% | -0.032 | 56A<sup>*</sup>, 73A<sup>**</sup>, 91A<sup>***</sup>, PS026<sup>**</sup> |
| JC2 | 3.500 | 2.002 | 0.711 | 0.358 | 0.374 | 0.101 | 0.358 | 2.959 | 11 | 79% | 0.063 | 56A<sup>**</sup>, 73A<sup>***</sup>, 91A<sup>*</sup>, AG8073<sup>**</sup>, Pae100<sup>**</sup> |
| JC3 | 4.250 | 2.779 | 0.934 | 0.408 | 0.454 | 0.055 | 0.408 | 3.802 | 12 | 80% | 0.127 | 56A<sup>***</sup>, 73A<sup>***</sup>, 91A<sup>***</sup>, P05<sup>**</sup>, Pae100<sup>***</sup> |
| JC4 | 3.438 | 1.924 | 0.733 | 0.425 | 0.399 | -0.022 | 0.425 | 2.953 | 11 | 85% | -0.04 | 56A<sup>***</sup>, 73A<sup>***</sup>, P10<sup>*</sup>, Pae100<sup>***</sup> |
| JC5 | 3.063 | 2.132 | 0.707 | 0.438 | 0.388 | -0.160 | 0.438 | 2.779 | 9 | 64% | -0.098 | 56A<sup>*</sup>, 73A<sup>***</sup>, PAG1<sup>*</sup> |
| M1 | 3.063 | 1.811 | 0.580 | 0.320 | 0.329 | 0.000 | 0.320 | 2.365 | 9 | 75% | 0.041 | 56A<sup>***</sup>, 73A<sup>***</sup>, 91A<sup>**</sup>, PAG1<sup>***</sup>, PS004<sup>***</sup> |
| M2 | 3.625 | 2.307 | 0.689 | 0.314 | 0.352 | 0.065 | 0.314 | 2.869 | 12 | 75% | 0.12 | 56A<sup>***</sup>, 73A<sup>***</sup>, 91A<sup>*</sup>, PAG1<sup>***</sup>, PS004<sup>***</sup>, PS026<sup>*</sup> |
| M3 | 3.500 | 2.584 | 0.767 | 0.385 | 0.398 | 0.079 | 0.385 | 3.087 | 14 | 78% | 0.055 | 56A<sup>***</sup>, 73A<sup>***</sup>, P03<sup>**</sup>, P05<sup>*</sup>, P10<sup>*</sup>, Pae100<sup>***</sup>, PS004<sup>**</sup>, PS026<sup>*</sup> |
| M4 | 2.563 | 1.859 | 0.593 | 0.325 | 0.335 | 0.071 | 0.325 | 2.563 | 9 | 69% | 0.083 | 56A<sup>*</sup>, 73A<sup>**</sup>, Pae100<sup>**</sup> |
| mean | 3.637 | 2.322 | 0.777 | 0.394 | 0.405 | 0.032 | 0.394 | 3.152 | 11.364 | 78% | 0.0512 |  |

$N_a$ : The observed number of allele.
$N_e$ : The effective number of alleles.
$I$ : Shannon's information index.
$H_o$ : Observed heterozygosity.
$H_e$ : Expected heterozygosity.
$F$ : Fixation index.
$PIC$ : Polymorphism information content.
$A_r$ : Allelic richness.
$NPL$ : the number of polymorphic loci.

### 3.3 Genetic differentiation and gene flow among populations

Genetic differentiation (Fst) among pairs of populations was highly significant (*P*<0.001), varying from 0.041 (between JC1 and JC2) to 0.234 (between DB2 and M2), with a mean value of 0.098 (*P*<0.001; Table 5) measured across eleven populations based on 16 markers. Conversely, the values for gene flow (Nm) between populations varied from 0.820 (between DB2 and M2) to 5.890 (between JC1 and JC2), with a mean value of 2.781 (Table S1). At the locus level, the genetic differentiation coefficient (Fst) and gene flow (Nm) calculated from F-statistics at each locus in the species were significantly different. Paired comparison of genetic differentiation between populations indicated that the maximum values of Fst and Nm were detected at locus 50F (0.430) and PSESP5 (2.735), respectively. The genetic differentiation coefficient (Fst) was estimated to be 0.193 for the 16 loci (ranging from 0.084 at locus PSESP5 to 0.430 at 50F) (Table 2), indicating that the self-crossing rate was very low at the species level and genetic differentiation was moderate (0.15<Fst<0.25). Additionally, the maximum differentiation coefficient (Fit) among individual plants occurred at locus 56A (0.496) (Table S2), with a mean of 0.220. The mean inbreeding coefficient (Fis) was 0.038.

**Table 5.** Genetic differentiation coefficient F_st_ between population. Note: F_st_ values are below the diagonal and associated P-values above.

|  | P-values above. |  |  |  |  |  |  |  |  |  |  |
| --- | --- | --- | --- | --- | --- | --- | --- | --- | --- | --- | --- |
|  | DB1 | DB2 | JC1 | JC2 | JC3 | JC4 | JC5 | M1 | M2 | M3 | M4 |
| DB1 | 0 | 0.000 | 0.000 | 0.000 | 0.000 | 0.000 | 0.000 | 0.000 | 0.000 | 0.000 | 0.000 |
| DB2 | 0.065 | 0 | 0.000 | 0.000 | 0.000 | 0.000 | 0.000 | 0.000 | 0.000 | 0.000 | 0.000 |
| JC1 | 0.094 | 0.154 | 0 | 0.000 | 0.000 | 0.000 | 0.000 | 0.000 | 0.000 | 0.000 | 0.000 |
| JC2 | 0.098 | 0.179 | **0.041** | 0 | 0.000 | 0.000 | 0.000 | 0.000 | 0.000 | 0.000 | 0.000 |
| JC3 | 0.095 | 0.161 | 0.063 | 0.052 | 0 | 0.000 | 0.000 | 0.000 | 0.000 | 0.000 | 0.000 |
| JC4 | 0.102 | 0.165 | 0.059 | 0.056 | 0.082 | 0 | 0.000 | 0.000 | 0.000 | 0.000 | 0.000 |
| JC5 | 0.102 | 0.209 | 0.072 | 0.063 | 0.101 | 0.08 | 0 | 0.000 | 0.000 | 0.000 | 0.000 |
| M1 | 0.1 | 0.16 | 0.073 | 0.07 | 0.083 | 0.106 | 0.094 | 0 | 0.000 | 0.000 | 0.000 |
| M2 | 0.136 | **0.234** | 0.076 | 0.081 | 0.08 | 0.105 | 0.088 | 0.109 | 0 | 0.000 | 0.000 |
| M3 | 0.096 | 0.165 | 0.054 | 0.064 | 0.059 | 0.101 | 0.099 | 0.061 | 0.047 | 0 | 0.000 |
| M4 | 0.134 | 0.22 | 0.078 | 0.071 | 0.068 | 0.102 | 0.098 | 0.085 | 0.044 | 0.043 | 0 |

Nei’s genetic distance calculated from a pairwise comparison varied from 0.058 (between M2 and M4) to 0.462 (between DB2 and M2) based on SSR markers, with a mean value of 0.178, the majority of pairwise genetic distances occurring over the range 0.1–0.3.

Genetic identity was also examined and ranged from 0.63 to 0.944 (mean 0.841) among the different populations, the maximum observed between M2 and M4 (Table 6). A Mantel test conducted for *P. decomposita* indicated significant positive correlation between geographic and genetic distance among populations (*r*=0.592, *P*<0.001) (Fig. 1), in line with the IBD (isolation by distance) model. Approximately 35% (*r*^2^=0.35) of genetic distance was a result of geographic distance among populations. Results of the AMOVA calculations demonstrated that 81.7% of the total molecular variation was due to differences within regions, while the remainder (18.3%) occurred among regions (*P*<0.001). At the population level, 73.48% of total variation resulted predominantly from individual differentiation within population, the remainder (only 26.52%) resulting from variation among populations (all *P*<0.001). When total variation was grouped into three hierarchical components, analysis by AMOVA revealed that the proportion of maximum variation (70.61%) was still brought about by genetic differentiation within populations (*P*<0.001), whereas 13.39% (*P* <0.001) and 16% (*P* <0.001) of total genetic variation resulted from genetic differentiation among regions and among populations within regions, respectively (Table 7). Therefore, significant differences in genetic differentiation existed among the 11 populations (Table 7).

**Fig. 1.**
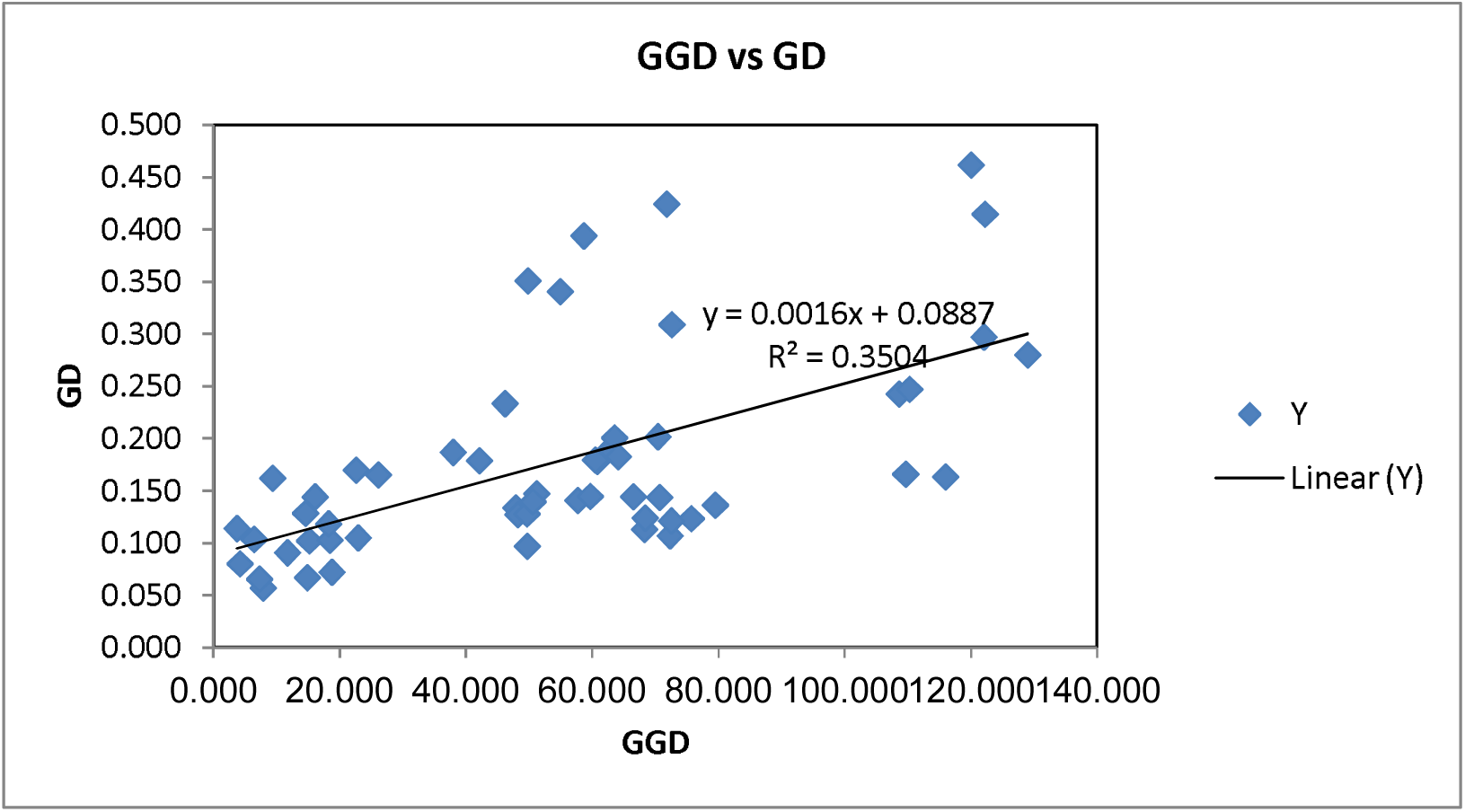
Correlation test of genetic distance (GD) and geographic distance (GGD)

**Table 6.** Nei’s genetic distances (below diagonal) and Nei’s genetic identity values (above diagonal) are given below for 11 populations. Bold character indicates highest Nei’s genetic distance between population DB2 and M2, while italic character displays the lowest genetic distance between population M2 and M4.

| Nei's Genetic Distance vs<br>Nei's Genetic Identity | DB1 | DB2 | JC1 | JC2 | JC3 | JC4 | JC5 | M1 | M2 | M3 | M4 |
| --- | --- | --- | --- | --- | --- | --- | --- | --- | --- | --- | --- |
| DB1 | – | 0.879 | 0.837 | 0.836 | 0.829 | 0.792 | 0.835 | 0.849 | 0.784 | 0.847 | 0.781 |
| DB2 | 0.129 | – | 0.734 | 0.711 | 0.704 | 0.674 | 0.654 | 0.756 | **0.63** | 0.743 | 0.66 |
| JC1 | 0.178 | 0.309 | – | 0.93 | 0.9 | 0.903 | 0.892 | 0.868 | 0.875 | 0.907 | 0.88 |
| JC2 | 0.179 | 0.341 | 0.073 | – | 0.923 | 0.901 | 0.902 | 0.884 | 0.865 | 0.893 | 0.883 |
| JC3 | 0.187 | 0.351 | 0.105 | 0.081 | – | 0.85 | 0.844 | 0.872 | 0.866 | 0.898 | 0.885 |
| JC4 | 0.234 | 0.394 | 0.102 | 0.104 | 0.163 | – | 0.865 | 0.817 | 0.827 | 0.818 | 0.833 |
| JC5 | 0.18 | 0.425 | 0.114 | 0.103 | 0.17 | 0.145 | – | 0.865 | 0.88 | 0.863 | 0.869 |
| M1 | 0.164 | 0.28 | 0.141 | 0.124 | 0.137 | 0.202 | 0.145 | – | 0.847 | 0.913 | 0.888 |
| M2 | 0.243 | **0.462** | 0.134 | 0.145 | 0.144 | 0.19 | 0.128 | 0.166 | – | 0.935 | **0.944** |
| M3 | 0.167 | 0.298 | 0.097 | 0.114 | 0.107 | 0.201 | 0.148 | 0.091 | 0.067 | – | 0.936 |
| M4 | 0.247 | 0.415 | 0.128 | 0.124 | 0.122 | 0.183 | 0.14 | 0.119 | *0.058* | 0.066 | – |

**Table 7.** Analysis of molecular variance (AMOVA) for 11 populations of *Paeonia decomposita*. Note: d.f. = degree of freedom; \*\*\**P* < 0.001.

| Source of variation | d.f. | Sum of squares | Variance components | Percentage of variation | *P*-value |
| --- | --- | --- | --- | --- | --- |
| Among regions | 2 | 348.45 | 2.02 | 18.30 | <0.001 |
| Within regions | 255 | 2299.61 | 9.02 | 81.70 | <0.001 |
| Among populations | 10 | 726.45 | 2.81 | 26.52 | <0.001 |
| Within populations | 247 | 1921.61 | 7.78 | 73.48 | <0.001 |
| Among regions | 2 | 348.45 | 1.48 | 13.39 | <0.001 |
| Among populations | 8 | 378.01 | 1.76 | 16.00 | <0.001 |
| within regions |  |  |  |  |  |
| Within populations | 247 | 1921.61 | 7.78 | 70.61 | <0.001 |

### 3.4 population structure and genetic relationship

The MCMC algorithm of reconstruction of SSR markers is displayed in Fig. 2. Bayesian analysis of population structure indicated that the optimal number of genetic clusters equaled 3 when ΔK was at its maximum for K=3 based on the method of Evanno *et al*. (2005) (Fig. 2), suggesting that this was the most likely number of genetic clusters. Thus, all 11 populations under study were split into three distinct genetic clusters (Fig. 3). Cluster 1 contained 49 individual plants collected from two populations in Danba county, Cluster 2 consisted of 97 individual plants sampled from five populations in Jinchuan county and the remaining 112 arising from four populations in Maerkang county were assigned to Cluster 3. It was clearly apparent that the three genetic clusters were identical to the clusters identified in PcoA, representing the natural distribution of *P. decomposita*. Principal coordinate analysis (PCoA) calculated from the genetic distance of 16 microsatellite primers between populations revealed a genetic structure that is presented in Fig. 4. The percentage variation attributable by the three principle coordinate axes was 76.66% (axis 1 – 50.71%, axis 2 – 16.95% and axis 3 – 9.00%). All populations were represented by the three groups.

**Fig. 2.**
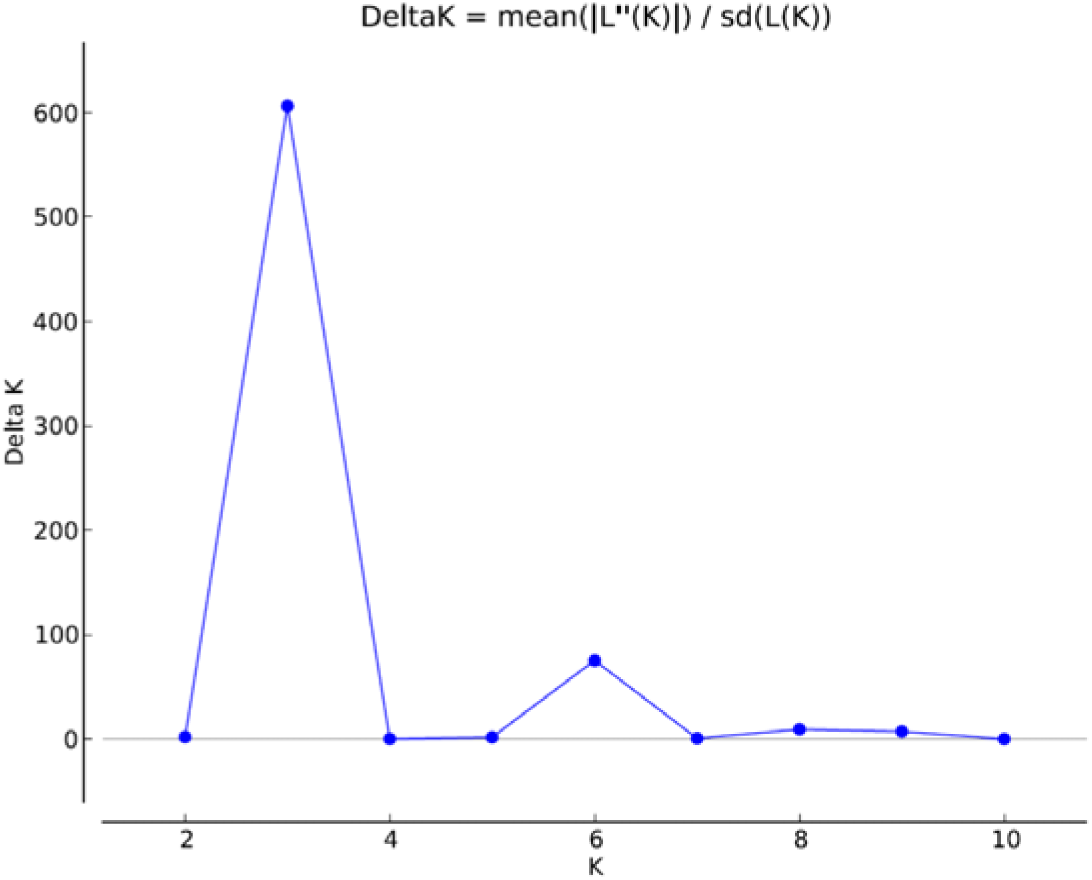
The distribution of ΔK over K=1–10

**Fig. 3.**
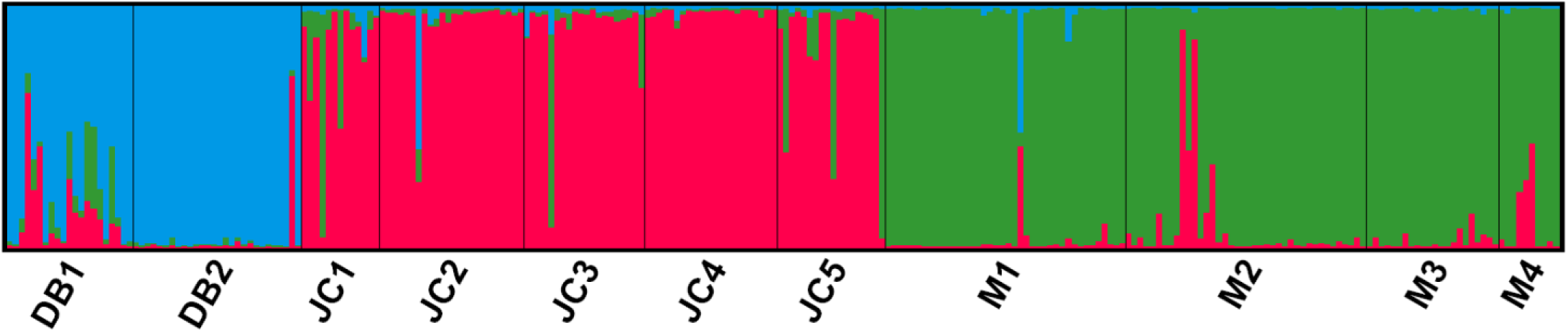
Genetic structure of 11 populations as inferred by STRUCTURE with SSR markers data set

**Fig. 4.**
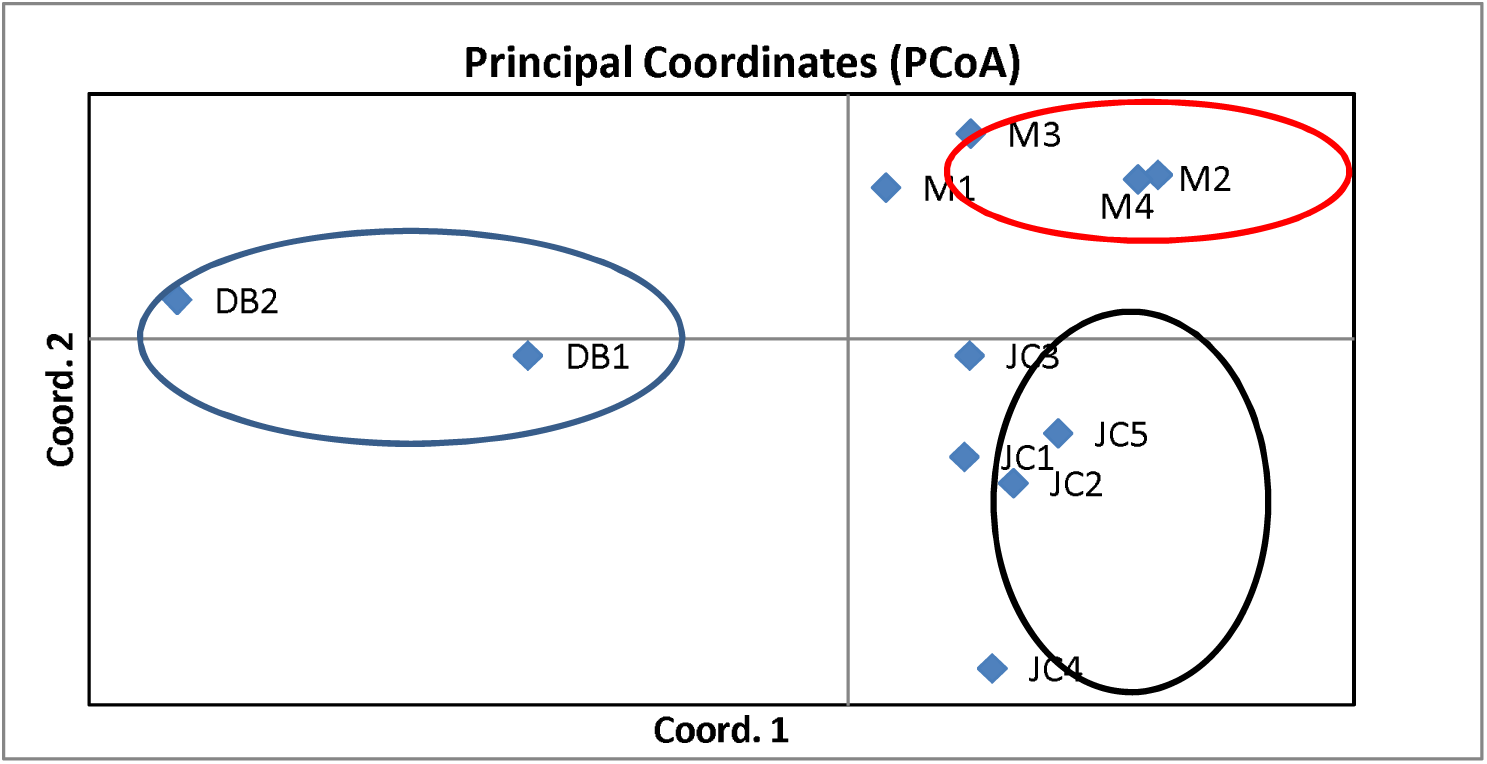
Principle Coordinate Analysis (PCoA) plot of the 11 populations showing three main clusters.

Group one included two populations from Danba county, group two principally comprised five populations from Jinchuan county and group three included four populations from Maerkang county. Furthermore, the results of PCoA was consistent with those of structure analysis and supported the UPGMA clustered tree, as follows.

The UPGMA dendrogram was constructed from Nei’s genetic distance values, accurately reflecting the genetic relationships among and within populations. The UPGMA tree indicated that the 11 populations could be divided into two major clades: 1 and 2 (Fig. 5).

**Fig. 5.**
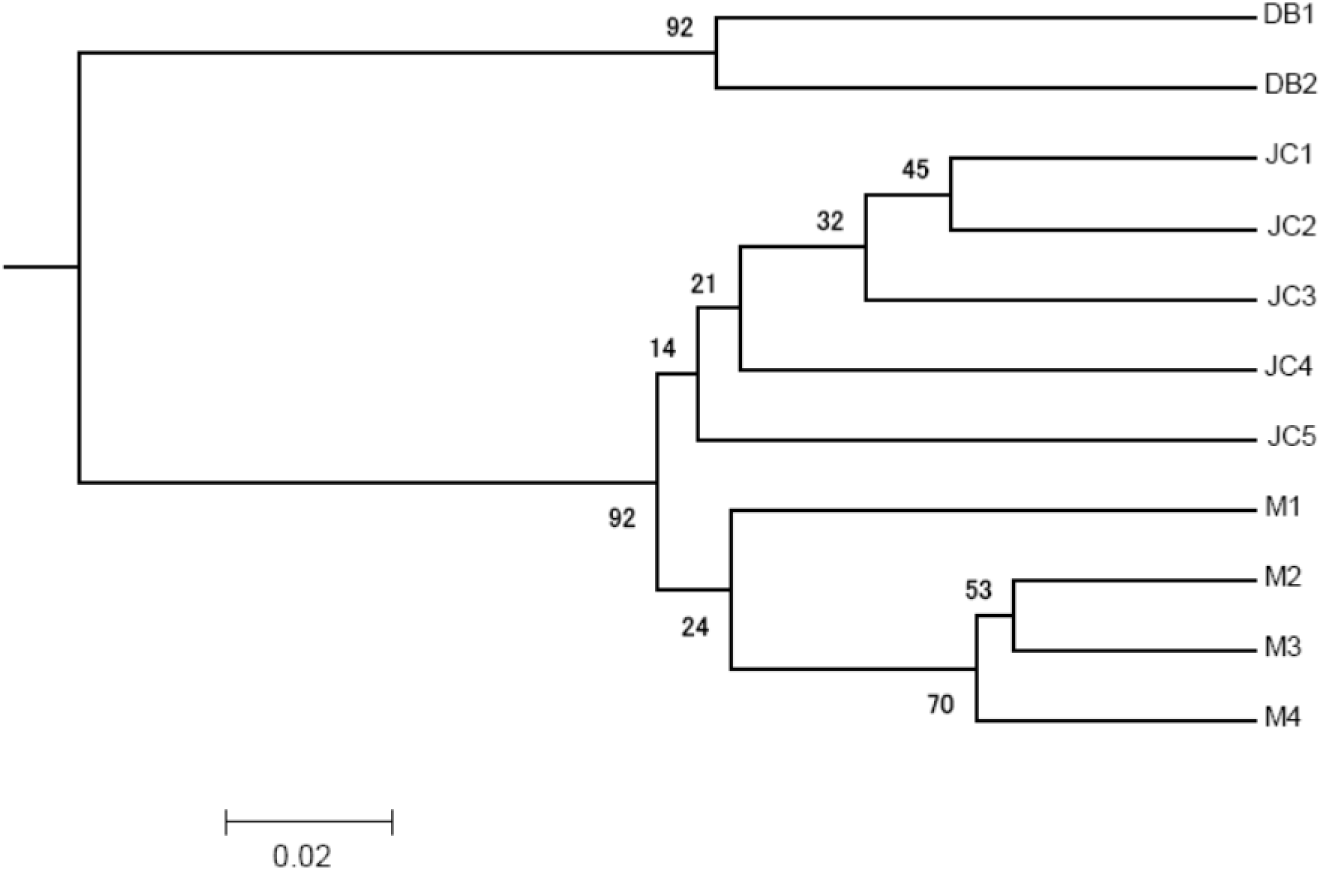
UPGMA dendrogram based on Nei’ genetic distance using SSR marker analysis. Branch length represents genetic distance, and the value on the branch is support rate.

Clade 1 included two populations, namely DB1 and DB2, with clade 2 consisting of the remaining 9 populations, which were further divided into two small branches: five populations (JC1, JC2, JC3, JC4 and JC5) from Jinchuan county formed one small branch and four (M1, M2, M3 and M4) from Maerkang county formed another. UPGMA also indicated that close genetic populations were distributed in geographic proximity to each other, for example, JC1/JC2 and M2/M3 were closely related and clustered together because they were closest in terms of genetic distance.

## 4. Discussion

In recent years, an increasing number of researchers have realized that it is important to maintain the genetic diversity of natural populations to ensure the continuing survival, fitness and potential for evolution of a species (Frankham *et al*., 2002). Traditionally, analysis of differences in plant morphology and physiological traits have been used to evaluate diversity. However, only limited information was available for this species using these methods, because such traits are not stable under different environmental conditions.

Recently, a range of DNA molecular marker techniques have been used to analyze tree peonies, including the use of RFLP (Zhao *et al*., 2004), RAPD (Zou *et al*., 1999; Su *et al*., 2006), ISSR (Suo *et al*., 2005) and AFLP markers (Liu *et al*., 2006). However, these studies were focused on investigating the phylogenetic relationships among interspecies or wild species and it is generally recognized that a greater number of molecular markers are required to conduct genetic studies of *Paeonia* species. SSR is the most practical molecular marker in studies of population genetics because it is able to measure codominant alleles and display high levels of polymorphism. The present study is the first to investigate the genetic diversity and population structure of *P. decomposita* through microsatellite markers, important for the conservation, management and greater understanding of its genetic relationships.

### 4.1 SSR marker polymorphism

Sixteen SSR markers amplified a total of 122 alleles, a mean of 7.625 alleles/locus, exhibiting relatively high polymorphism. Mean observed heterozygosity (Ho=0.385) was relatively important, revealing high heterozygosity within individuals. The majority of primers (11/16) identified a moderate and low level of polymorphism. He was higher than Ho in 15 of the 16 SSR loci except for 73A, which exhibited considerable differences between loci in the fixation index (F) (range: –0.798∼0.389, mean: 0.047).

In natural populations, high PIC values represent high genetic differentiation, in turn implying a complex genetic background at the molecular level. According to the standards described by Botstein *et al*. (1980), the majority of primers had moderate or low PIC, 5 of the 16 primer pairs generating low polymorphic loci (0<PIC<0.25), 4 that were moderate (0.25<PIC<0.50), and 7 with highly polymorphic loci (0.50<PIC<1.00). High PIC values may be due to a greater number of repeat motifs observed in the primers in this study. Thus, SSR markers were helpful for evaluation of genetic diversity among populations and for investigating population structure.

In general, higher genetic diversity existed within populations with less genetic differentiation existing among populations in outcrossing and woody plants (Hamrick and Godt, 1996). In this study, the 16 SSR primers demonstrated a higher genetic diversity level (He*=*0.492) and lower genetic differentiation (Fst*=*0.193) than those in the ISSR study (Tong *et al*., 2016).

### 4.2 Genetic diversity of *P. decomposita* among populations

Differences in genetic diversity may result from a small number of factors, for example, life-history traits or geographic traits of a species (Nybom, 2004). In general, less genetic diversity exists in an endemic species that is not widely distributed compared with that found in a widespread species (Hamrick and Godt, 1996; Huh and Huh, 1999), usually because their population numbers are limited, and as they are isolated from other populations they adapt to their particular habitat (Barrett and Kohn, 1991).

Our study demonstrated that the genetic diversity level of *P. decomposita* was moderate (Ho = 0.394, He = 0.405) among *Paeonia* species even though it is a rare and endangered endemic species. Compared with previous research of wild tree peonies, the genetic diversity parameters observed in this study were slightly lower than those of *P. jishanensis* (Ho = 0.446) (Xu *et al*., 2016) and *P. rockii* (Ho = 0.459, He = 0.492) (Yuan *et al*., 2012), but higher than those of *P. ostii* (Ho = 0.343, He = 0.321) (Peng *et al*., 2017), *P. jishanensis* (He = 0.340) (Xu *et al*., 2016), *P. delavayi* (Ho = 0.334, He = 0.369) (Zhang *et al*., 2018) and *P. ludlowii* (Ho = 0.014, He = 0.013) (Zhang *et al*., 2018). The genetic diversity analysis using ISSR markers also indicated a level for *P. decomposita* that was not high (Tong *et al*., 2016) and similar to our results.

Levels of genetic diversity in *P. decomposita* (He = 0.405) were lower than both “endemic” species (He=0.420) and “widespread” species (He=0.620) (Nybom, 2004). This result was obtained for three potential reasons. Firstly, *P. decomposita* is a long-lived shrub that may exhibit genetic diversity from ancestral populations (Luan *et al*., 2006; Setoguchi *et al*., 2011). Secondly, the mating system is often viewed as a principal factor influencing the genetic diversity of species. Thirdly, the sporadic and narrow distribution range, the small size of populations and large spatial distances between populations limited pollination among populations.

Current methods of analysis have considerably improved our understanding of genetic diversity in populations of *P. decomposita*, in which the polymorphism levels varied between populations. In this study, genetic diversity (I, Ho, He, PIC) at a population level was relatively uniform and high in populations DB1 and DB2, related to low levels of human disturbance and a large population size in Danba. Therefore, Danba represents the major genetic diversity center of the species. Mean expected heterozygosity was lower than mean observed heterozygosity in populations JC1, JC4 and JC5, *i.e.*, there were excessive numbers of heterozygotes. Estimation of fixation index (F) revealed that three populations (JC1, JC4, JC5: negative values) displayed an excess of heterozygotes indicating outbreeding and eight other populations (positive value) had an excess of homozygotes associated with inbreeding (Table 4). Mean positive inbreeding coefficient (Fis) values (0.051) indicated an excess of homozygotes in *P. decomposita*, consistent with the self-compatibility system of this species (Table 3).

The results strengthened the assumption that endangered plants with a narrow distribution are generally aplastic. A reduction in genetic variation might suggest a decline in adaptation to a changing environment, leading to increased danger of extinction and increased inbreeding (Tansley and Brown, 2000; Frankham *et al*., 2002; Frankham *et al*., 2010).

### 4.3 Genetic differentiation and gene flow

Two important parameters, gene flow and the genetic differentiation coefficient, are employed to assess the genetic structure of a population (Hamrick and Godt, 1990). Gene flow and the genetic differentiation coefficient are negatively correlated (Grant, 1986).

Gene flow is a basic micro-evolutionary phenomenon that destroys genetic differentiation among populations and affects maintenance of genetic diversity (Slatkin, 1994; Yao *et al*., 2007). Many endangered plants currently occur only as highly isolated and narrowly distributed within a few small populations, possibly leftovers of a formerly widespread species, which had large and continuous populations (Yao *et al*., 2007; Setoguchi *et al*., 2011). In the present research study, gene flow (mean Nm value) among *P. decomposita* populations was >1 and did not exhibit genetic differentiation, resulting from genetic drift (Hamrick *et al*., 1992). Genetic drift has not yet become a predominant factor influencing the genetic structure of *P. decomposita*. However, *P. decomposita* populations are now affected by fragmentation and vandalism, with genetic exchanges frequently occurring within most populations which may become continuously distributed. These causes, together with the fact that natural populations are spatially distant (isolated by mountain and river barriers), may result in genetic drift occurring gradually.

Though diversity has mostly occurred within populations, the majority of genetic differentiation among populations has occurred at a moderate and low level except the high levels of genetic differentiation between DB2 and other populations (Table 5), according to the scale suggested by Wright (1978). Mean Fst among the 11 populations indicates relatively moderate but significant overall genetic differentiation among the populations of the species. The Fst value observed among the 11 populations might result from isolation of the populations, geographic distance and environmental adaptation.

The AMOVA results (*P*<0.001) also support population differentiation. AMOVA revealed the presence of significant variation among and within the populations, with considerable genetic diversity within rather than among populations, a situation identical to that observed with other cross-pollinating species in *Paeonia* (Yuan *et al*., 2012; Peng *et al*., 2017; Zhang *et al*., 2018) and other studies exploring ISSR markers (Tong *et al*., 2016).

Outcrossing and long-lived plants demonstrated that the majority of their genetic variation exists within populations, while selfing plants maintain the majority of the genetic variation among populations (Nybom, 2004).

The genetic differentiation level observed in the present study was lower than that reported by a previous study by Tong *et al*., in which 32.57% of genetic variation existed among the populations and 67.43% within the populations (Tong *et al*., 2016), possibly due to differences in population numbers studied, or molecular markers investigated.

### 4.4 Population structure and genetic relationship

Increasing numbers of methods are being used to detect genetic diversity and population structure (Zhang *et al*., 2006; Zong *et al*., 2009; Tan *et al*., 2011; Zhang *et al*., 2012; Lai *et al*., 2014). It is advisable to combine three effective techniques and so we consider that the combination of PCoA, Structure and UPGMA analysis is able to produce reliable results.

UPGMA was able to expound intuitive relationships although it cannot fully categorize populations. Conversely, Structure software can objectively categorize populations and produce plans for breeding. Therefore, this method was regarded as the most suitable to categorize populations.

There was a clear genetic structure among the *P. decomposita* populations, and the proportion exhibiting genetic differentiation among populations was 19.3% (F_st_*=*0.193). This result was supported by AMOVA analysis, in which the divided genetic variation was 26.52%.

In the present study, UPGMA cluster analysis grouped 11 populations collected from three different regions into two clades (Fig. 5), demonstrating that there were two distinct genetic groups in these areas. Analysis using Structure based on a Bayesian model indicated a maximum △K value when K=3, illustrating that the 11 populations were divided into three clusters, as shown in Fig. 4. It was clear that despite the occurrence of introgression, there were three distinct clusters. This suggests that analyses by Structure software were reliable. Furthermore, the PCoA results were identical to those from Structure and inconsistent with the UPGMA clustered tree.

In addition, the genetic relationships among populations reflected those populations’ natural geographical locations which were supported by an IBD (isolation-by-distance) model constructed using a Mantel test. This IBD model for *P. decomposita* indicated a significantly positive correlation (r=0.592, *P*<0.001) between geographic distance and genetic distance between populations. The closer the populations were geographically, the lower the genetic differentiation. Thus, the genetic differentiation among populations increased as distance among populations increased, and so Mantel test analysis suggested that the genetic clusters were significantly related to the populations’ geographic origins. The differences in genetic differentiation were due to geographic barriers, which isolated different gene pools. Inefficient pollen flow, close seed dispersal and low germination rates are latent reasons, which have led to three distinct *P. decomposita* gene pools.

Genetic distance is highly significant in every population relationship study. In general, the closer two individuals are, the higher the probability of a common ancestry. The Mantel test supported the UPGMA dendrogram constructed on genetic distance.

### 4.5 Conservation of populations *in situ* and *ex situ*

It is essential to understand the genetic diversity of a population, its structure and gene flow in order to create an appropriate management and conservation strategy. The population resource employed for reintroduction, including reproduction material and collection of germplasm must be optimal in terms of genetic variation, with low levels of inbreeding that restrains the growth of a population.

The management of collections and conservation of genetic resource must guarantee that the majority of existing variation is conserved. Conservation of diversity among populations must concentrate on maintaining the most genetically distinctive populations while conservation of diversity within populations must conserve large core populations, in which diversity is not lost due to genetic drift (Namkoong, 1988). In the case of *P. decomposita*, conservation must consider not only geographic distance between populations, but also the existence of different clusters and their different growth habitats. In every cluster, the priorities for the conservation of populations must be selected, by considering the level of genetic diversity, the state of a populations’ regeneration and its level of threat. Construction of big reserves with several populations in every cluster could guarantee a sample of gene pool, which could embrace the uniqueness and diversity that exists in all populations.

Genetic diversity is especially important in a species in order to preserve the latent evolutionary capacity to deal with changing environments. The maintenance of genetic diversity and evolutionary potential is a primary goal for the conservation of endangered species in management programs (Margules and Pressey, 2000; Frankham *et al*., 2002; Rodrigues *et al*., 2013). Therefore, information about genetic variation within and among populations in endangered and rare plants plays an important role in the process of formulating conservation and management strategies (Milligan *et al*., 1994). Thus, we suggest that three natural distribution areas should correspond to three management units. In view of the current circumstances in which a rapid fall in the numbers of populations and the extreme endangerment of their natural habitats, *in situ* and *ex situ* conservation are imperative. All populations, particularly those with high genetic levels of diversity or those with a large genetic difference, should be protected. *In situ* conservation is considered to be the most effective method of protecting endangered plants, through which the whole gene pool can be protected in a natural habitat. Small populations are more likely to become extinct due to habitat damage and environmental fluctuation. It is essential to conserve all individual plants and populations *in situ* for the sake of preserving genetic variation as far as possible.

Traditional methods of protection that primarily concentrate on *in situ* conservation, such as improving regeneration, controlling overgrazing and protecting natural habitats, may be sufficient to maintain the size of population. Consequently, it is essential to prevent a populations’ genetic homogeneity. *In situ* conservation must be introduced promptly by defining and introducing conservation reserves in core distribution regions and strictly prohibiting the harvesting of wild plants of *P. decomposita*. Populations with higher genetic diversity, for example, DB1 and DB2, must be given priority for conservation *in situ*. Much previous research has demonstrated that heterozygosity is the best way to ensure a populations’ fitness and potential for adaptation (Reed and Frankham, 2003). However, notable heterozygote deficit was found to exist in the majority of the populations tested, possibly a result of inbreeding in fragments of populations.

The populations are facing problems of habitat destruction, loss or fragmentation as a result of grazing (M2, JC5, D1, D2), over harvesting (M2, M3), abusive seed collection (JC2-5), being near villages, farm fields, orchards (JC2-5), or areas practically destroyed by urban expansion (M2, M4). Given this challenge, in addition to *in situ* conservation, it is very much advised that gene banks in both field and laboratory are established *ex situ* for each population for which protection is required for endangered plants (Heywood and Iriondo, 2003). Populations with high genetic diversity, for example, DB1 and DB2, must be concrete goals for *ex situ* conservation. Because the degree of genetic differentiation was low among populations, each may represent a large component of genetic variation in a species. Thus, seed collection tactics could be devised for the construction of an *ex situ* seed germplasm resource bank to collect as many samples of each population as possible from the whole natural geographical distribution with different genetic clusters, and conserve the germplasm using plant tissue culture techniques. In the course of *ex situ* conservation, artificial hybridization must be performed among populations with large genetic differences to rapidly improve heterozygosity. After *ex situ* cultivation of seeds collected from the field, saplings should be introduced into source sites. To summarize, *in situ* and *ex situ* conservation methods should be combined to protect valuable genetic resources.

## 5. Conclusions

Genetic information from this detailed study has provided first-hand data of the genetic diversity and structure of *P. decomposita* populations in the main distribution areas which are beneficial for developing measures to conserve and manage endangered and endemic plants. Among 11 populations from 11 sites across the majority of the distribution areas, 122 alleles were obtained in total with an average of 7.625 alleles per locus. Natural populations maintained moderate to low genetic diversity levels, high gene flow and low genetic differentiation among populations. AMOVA demonstrated that major variation existed within populations. From the results of Structure analysis, 11 natural populations were categorized into three groups by PCA cluster analysis, which should possibly be considered as three management units for the objective of conservation. These populations are a precious genetic resource for a future breeding plan and conservation strategy. The largest number of populations should be saved by *in situ* and *ex situ* conservation measures, taking precedence over those with genetic diversity and differentiation. This is the first time that the genetic diversity of *P. decomposita* has been studied using SSR, the results representing a reference for improving the germplasm and parental selection for breeding strategy plans.

In this study, the markers used allowed investigation of population structure, genetic diversity and proposed germplasm collection and a conservation strategy for *P. decomposita*. Important information about genetic structure was provided by these markers, which significantly contribute to future improvements and breeding plans for the species. The genetic diversity, population structure and genetic relationships between the populations through SSR analysis will be helpful for crop breeding, germplasm management and conservation. To conclude, these results provide value as an important resource to study genetic diversity and assist conservation and research plans in the future.

## Acknowledgements

This study was supported by the National Natural Science Foundation of China (grant nos. 31670345, 31860085)

---

**Table S1.** Pairwise population Fst values and estimates of Nm for 11 populations of *Paeonia decomposita*

| Pop1 | Pop2 | Fst | Nm | #Pop1 | #Pop2 |
| --- | --- | --- | --- | --- | --- |
| JC1 | JC2 | 0.041 | 5.89 | 13 | 24 |
| M3 | M4 | 0.043 | 5.592 | 22 | 10 |
| M2 | M4 | 0.044 | 5.493 | 40 | 10 |
| M2 | M3 | 0.047 | 5.108 | 40 | 22 |
| JC2 | JC3 | 0.052 | 4.516 | 24 | 20 |
| JC1 | M3 | 0.054 | 4.42 | 13 | 22 |
| JC2 | JC4 | 0.056 | 4.183 | 24 | 22 |
| JC1 | JC4 | 0.059 | 3.981 | 13 | 22 |
| JC3 | M3 | 0.059 | 3.964 | 20 | 22 |
| M1 | M3 | 0.061 | 3.816 | 40 | 22 |
| JC2 | JC5 | 0.063 | 3.726 | 24 | 18 |
| JC1 | JC3 | 0.063 | 3.71 | 13 | 20 |
| JC2 | M3 | 0.064 | 3.64 | 24 | 22 |
| DB1 | DB2 | 0.065 | 3.606 | 21 | 28 |
| JC3 | M4 | 0.068 | 3.443 | 20 | 10 |
| JC2 | M1 | 0.07 | 3.319 | 24 | 40 |
| JC2 | M4 | 0.071 | 3.255 | 24 | 10 |
| JC1 | JC5 | 0.072 | 3.238 | 13 | 18 |
| JC1 | M1 | 0.073 | 3.153 | 13 | 40 |
| JC1 | M2 | 0.076 | 3.033 | 13 | 40 |
| JC1 | M4 | 0.078 | 2.941 | 13 | 10 |
| JC4 | JC5 | 0.08 | 2.893 | 22 | 18 |
| JC3 | M2 | 0.08 | 2.861 | 20 | 40 |
| JC2 | M2 | 0.081 | 2.818 | 24 | 40 |
| JC3 | JC4 | 0.082 | 2.814 | 20 | 22 |
| JC3 | M1 | 0.083 | 2.752 | 20 | 40 |
| M1 | M4 | 0.085 | 2.691 | 40 | 10 |
| JC5 | M2 | 0.088 | 2.577 | 18 | 40 |
| JC5 | M1 | 0.094 | 2.41 | 18 | 40 |
| DB1 | JC1 | 0.094 | 2.404 | 21 | 13 |
| DB1 | JC3 | 0.095 | 2.38 | 21 | 20 |
| DB1 | M3 | 0.096 | 2.352 | 21 | 22 |
| DB1 | JC2 | 0.098 | 2.31 | 21 | 24 |
| JC5 | M4 | 0.098 | 2.296 | 18 | 10 |
| JC5 | M3 | 0.099 | 2.27 | 18 | 22 |
| DB1 | M1 | 0.1 | 2.238 | 21 | 40 |
| JC3 | JC5 | 0.101 | 2.223 | 20 | 18 |
| JC4 | M3 | 0.101 | 2.22 | 22 | 22 |
| DB1 | JC4 | 0.102 | 2.213 | 21 | 22 |
| DB1 | JC5 | 0.102 | 2.204 | 21 | 18 |
| JC4 | M4 | 0.102 | 2.195 | 22 | 10 |
| JC4 | M2 | 0.105 | 2.139 | 22 | 40 |
| JC4 | M1 | 0.106 | 2.107 | 22 | 40 |
| M1 | M2 | 0.109 | 2.043 | 40 | 40 |
| DB1 | M4 | 0.134 | 1.609 | 21 | 10 |
| DB1 | M2 | 0.136 | 1.588 | 21 | 40 |
| DB2 | JC1 | 0.154 | 1.372 | 28 | 13 |
| DB2 | M1 | 0.16 | 1.313 | 28 | 40 |
| DB2 | JC3 | 0.161 | 1.304 | 28 | 20 |
| DB2 | M3 | 0.165 | 1.267 | 28 | 22 |
| DB2 | JC4 | 0.165 | 1.264 | 28 | 22 |
| DB2 | JC2 | 0.179 | 1.143 | 28 | 24 |
| DB2 | JC5 | 0.209 | 0.947 | 28 | 18 |
| DB2 | M4 | 0.22 | 0.887 | 28 | 10 |
| DB2 | M2 | 0.234 | 0.82 | 28 | 40 |
| Mean |  | 0.098 | 2.781 |  |  |

**Table S2.** F-statistics and estimates of Nm over all pops for each locus of *Paeonia decomposita*

| Locus | Fis | Fit | Fst | Nm |
| --- | --- | --- | --- | --- |
| 56A | 0.374 | 0.496 | 0.195 | 1.033 |
| Pae100 | 0.284 | 0.452 | 0.235 | 0.815 |
| 50F | -0.026 | 0.416 | 0.430 | 0.331 |
| AG8073 | 0.148 | 0.405 | 0.302 | 0.577 |
| WD09 | 0.245 | 0.404 | 0.211 | 0.936 |
| 91A | 0.308 | 0.404 | 0.138 | 1.556 |
| PAG1 | 0.154 | 0.283 | 0.153 | 1.385 |
| PS026 | 0.020 | 0.265 | 0.250 | 0.751 |
| PO5 | 0.081 | 0.242 | 0.175 | 1.175 |
| PSMP2 | -0.002 | 0.150 | 0.151 | 1.400 |
| P10 | -0.001 | 0.145 | 0.146 | 1.467 |
| PSESP5 | 0.062 | 0.141 | 0.084 | 2.735 |
| P03 | -0.182 | 0.119 | 0.255 | 0.732 |
| PS004 | -0.031 | 0.112 | 0.139 | 1.554 |
| P12 | -0.035 | 0.108 | 0.139 | 1.552 |
| 73A | -0.784 | -0.622 | 0.091 | 2.502 |
| Mean | 0.038 | 0.220 | 0.193 | 1.281 |
| SE | 0.066 | 0.066 | 0.022 | 0.162 |

## References

Aboukhalid K, Machon N, Lambourdière J, Abdelkrim J, Bakha M, Douaik A, Korbecka-Glinka G, Gaboun F, Tomi F, Lamiri A, AlFaiz C. 2017. Analysis of genetic diversity and population structure of the endangered *Origanum compactum* from Morocco, using SSR markers: Implication for Conservation. Biological Conservation 212, 172–182. http://dx.doi.org/10.1016/j.biocon.2017.05.030

Agarwal M, Shrivastava N, Padh H. 2008. Advances in molecular marker techniques and their applications in plant sciences. Plant Cell Reports 27, 617–631. https://doi.org/10.1007/s00299-008-0507-z

Barrett, SCH, Kohn JR. 1991. Genetic and evolutionary consequences of small population size in plants: implications for conservation. In: Falk DA, Holsinger KE (eds) Genetics and Conservation of Rare Plants. New York: Oxford University Press, 3–30.

Botstein D, White RL, Skolnick M, Davis RW. 1980. Construction of a genetic linkage map in man using restriction fragment length polymorphisms. American Journal of Human Genetics 32, 314–331.

Cai CF. 2015. High-density genetic linkage map construction and QTLs analyses for phenofypic traits in tree peony. Dissertation, Beijing Forestry University.

Cheng FY. 2007. Advances in the breeding of tree peonies and a cultivar system for the cultivar group. International Journal of Plant Science 1, 89–104.

Cheng FY, Li JJ, Chen DZ. 1997. The natural propagation characteristics of wild tree peony species in China. Acta Horticultural Sinica 24, 180–184.

Cohen JI, Williams JT, Plucknett DL, Shands H. 1991. Ex situ conservation of plant genetic resources: global development and environmental concerns. Science 253, 866–872. https://doi.org/10.1126/science.253.5022.866

Earl DA, vonHoldt BM. 2011. STRUCTURE HARVESTER: a website and program for visualizing STRUCTURE output and implementing the Evanno method. Conservation Genetics and Resources 4, 359–361. http://dx.doi.org/10.1007/s12686-011-9548-7

Evanno G, Regnaut S, Goudet J. 2005. Detecting the number of clusters of individuals using the software STRUCTURE: a simulation study. Molecular Ecology 14, 2611–2620. http://dx.doi.org/10.1111/j.1365-294X.2005.02553.x

Felsenstein J. 2005. PHYLIP (Phylogeny Inference Package) Version3.6.7. Department of Genome Sciences, University of Washington, Seattle, WA, USA.

Frankham R, Ballou JD, Briscoe DA. 2002. Introduction to Conservation Genetics. Cambridge University Press, Cambridge, UK.

Frankham R, Ballou JD, Briscoe DA. 2010. Introduction to Conservation Genetics. Cambridge University Press, Cambridge, UK.

Gao ZM, Wu J, Liu ZA, Wang LS. Ren HX, Shu QY. 2013. Rapid microsatellite development for tree peony and its implications. BMC Genomics 14, 886–896. http://dx.doi.org/10.1186/1471-2164-14-886

Gilmore B, Bassil N, Nyberg A, Knaus B, Smith D, Barney DL, Hummer K. 2013. Microsatellite marker development in peony using next generation sequencing. Journal of the American Society for Horticultural Science 138, 64–74. http://dx.doi.org/10.21273/JASHS.138.1.64

Goudet J. 2001. FSTAT, a program to estimate and test gene diversities and fixation indices (version 2.9.3). Lausanne: Université de Lausanne, Disponívelem: http://www.unil.ch/izea/softwares/fstat.html (accessed 22.10.2001).

Grant V. 1986. The evolutionary process: a critical study of evolutionary theory. Studies in History and Philosophy of Science 17, 65–98.

Hamrick JL, Godt MJW. 1990. Allozyme diversity in plant species. In: Brown AHD, Clegg MT, Kahler AL, Weir BS (eds). Plant Population Genetics, Breeding and Genetic Resources. Sinauer Associates, Sunderland, MA, 43–63.

Hamrick JL, Godt MJW. 1996. Effects of life history traits on genetic diversity in plant species. Philosophical transactions of the Royal Society B-Biological Sciences 351, 1291–1298.

Hamrick JL, Godt MW, Sherman-Broyles SL. 1992. Factors influencing levels of genetic diversity in woody plant species. New Forests 6, 95–124. http://dx.doi.org/10.1007/BF00120641

Han JG, Li XQ, Liu Z, Hu YH. 2014. Potential applications of tree peony as an oil plant. Grain and Oil 27, 21–25.

Heywood VH, Iriondo JM. 2003. Plant conservation: old problems, new perspectives. Biological Conservation 113, 321–335. http://dx.doi.org/10.1016/S0006-3207(03)00121-6

Homolka A, Berenyi M, Burg K, Kopecky D, Fluch S. 2010. Microsatellite markers in the tree peony, *Paeonia Suffruticosa* (Paeoniaceae). American Journal of Botany 97, e42–e44. http://dx.doi.org/10.3732/ajb.1000127

Hong DY. 1997. Notes on *Paeonia decomposita* Hand.-Mazz. Kew Bulletin 52, 957–963. http://dx.doi.org/10.2307/4117822

Hong DY. 2010. Peonies of the world: taxonomy and phytogeography, Royal Botanical Gardens Kew Publishing, Kew & Missouri Botanical Garden Press, St. Louis, London.

Hong DY. 2011. *Paeonia rotundiloba* (DY Hong) DY Hong: A new status in tree peonies (Paeoniaceae). Journal of Systematics and Evolution 49, 464–467. https://doi.org/10.1111/j.1759-6831.2011.00149.x

Hong DY, Pan KY. 1999. Taxonomical history and revision of *Paeonia* sect. Moutan (Paeoniaceae). Acta Phytotaxonomica Sinica 37, 351–368.

Hong DY, Pan KY, Turland JN. 2001. Paeoniaceae, Flora of China. Beijing: Science Press and Missouri Botanical Garden Press.

Hong DY, Zhou SL, He XJ, Yuan JH, Zhang YL, Cheng FY, Zeng L, Wang Y, Zhang XX. 2017. Current status of wild tree peony species with special reference to conservation. Biodiversity Science 25, 781–793. https://doi.org/10.17520/biods.2017129

Hou XG, Guo DL, Cheng SP, Zhang JY. 2011. Development of thirty new polymorphic microsatellite primers for P*aeonia suffruticosa*. Biological plantarum 55, 708–710. http://dx.doi.org/10.1007/s10535-011-0172-x

Hou XG, Guo DL, Wang J. 2011. Development and characterization of EST-SSR markers in *Paeonia suffruticosa* (Paeoniaceae). American Journal of Botany 98, e303–e305. http://dx.doi.org/10.3732/ajb.1100172

Huh M, Huh HW. 1999. Patterns of genetic diversity and population structure of the clonal herb, Potentilla fragarioides var. sprengeliana (Rosaceae) in Korea. Acta Botanica Sinica 42, 64–70.

Kalinowski ST, Taper ML, Marshall TC. 2010. Revising how the computer program CERVUS accommodates genotyping error increases success in paternity assignment. Molecular Ecology 19, 1512–1519. http://dx.doi.org/10.1111/j.1365-294X.2010.04544.x

Kumar GS, Singh R, Choudhury DR, Bharadwaj J, Gupta V, Singoded A. 2014. Genetic diversity and population structure study of drumstick (*Moringa oleifera* Lam.) using morphological and SSR markers. Industrial Crops and Products 60, 316–325. http://dx.doi.org/10.1016/j.indcrop.2014.06.033

Lai GT, Lai ZX, Liu WH, Ye W, Lin YL, Liu SC, Chen YK, Zhang ZH, Wu GJ. 2014. ISSR analysis of 3 natural populations of the wild banana distributed in the middle of Fujian province based on NTSYS and STRUCTURE. Chinese Journal of Tropical Crops 35, 223–231.

Li K, Zhou N, Li HY. 2012. Composition and function research of peony flowers and peony seeds. Food Research and Development 33, 228–230.

Lin QB, Zhou ZQ, Zhao X, Pan KY, Hong DY. 2004. Interspecific relationships among the wild species of *Paeonia* Section *Moutan* DC based on DNA sequences of Adh gene family. Acta Horticulturae Sinica 31, 627–632.

Litkowiec M, Lewandowski A, Wachowiak W. 2018. Genetic variation in *Taxus baccata* L.: A case study supporting Poland’s protection and restoration program. Forest Ecology and Management 409, 148–160. http://dx.doi.org/10.1016/j.foreco.2017.11.026

Liu P, Wang ZC, Shang F. 2006. AFLP analysis of genetic diversity of *Paeonia suffruticosa* cultivars in Henan Province. Acta Horticultural Sinica 33, 1369–1372.

Luan SS, Chiang TY, Gong X. 2006. High genetic diversity vs. low genetic differentiation in *Nouelia insignis* (Asteraceae), a narrowly distributed and endemic species in China, revealed by ISSR fingerprinting. Annals of Botany 98, 583–589. http://dx.doi.org/10.1093/aob/mcl129

Margules CR, Pressey RL. 2000. Systematic conservation planning. Nature 405, 243–253. http://dx.doi.org/10.1038/35012251

Meng L, Zheng GS. 2004. Phylogenetic relationship analysis among Chinese wild species and cultivars of *Paeonia* section *Moutan* using RAPD markers. Scientia Silvae Sinica 40, 110–115.

Milligan B, Leebens-Mack J, Strand A. 1994. Conservation genetics: beyond the maintenance of marker diversity. Molecular Ecology 3, 423–435. http://dx.doi.org/10.1111/j.1365-294X.1994.tb00082.x

Namkoong G. 1988. Sampling for germplasm collections. HortScience 23, 79–81.

Ni JL, Zhu AG, Wang XF, Xu Y, Sun ZM, Chen JH, Luan MB. 2018. Genetic diversity and population structure of ramie *(Boehmeria nivea* L.). Industrial Crops and Products 115, 340–347. http://dx.doi.org/10.1016/j.indcrop.2018.01.038

Nybom H. 2004. Comparison of different nuclear DNA markers for estimating intraspecific genetic diversity in plants. Molecular Ecology 13, 1143–1155. http://dx.doi.org/10.1111/j.1365-294X.2004.02141.x

Ouborg NJ. 2010. Integrating population genetics and conservation biology in the era of genomics. Biology Letters 6, 3–6. http://dx.doi.org/10.1098/rsbl.2009.0590

Peakall R, Smouse PE. 2012. GenAlEx 6.5, genetic analysis in Excel. Population genetic software for teaching and research – an update. Bioinformatics 28, 2537–2539. http://dx.doi.org/10.1093/bioinformatics/bts460

Peng LP, Cai CF, Zhong Y, Xu XX, Xian HL, Cheng FY, Mao JF. 2017. Genetic analyses reveal independent domestication origins of the emerging oil crop *Paeonia ostii*, a tree peony with a long-term cultivation history. Scientific Reports 7, 5340–5352. http://dx.doi.org/10.1038/s41598-017-04744-z

Pritchard JK, Stephens M, Donnelly P. 2000. Inference of population structure using multilocus genotype data. Genetics 155, 945–959.

Reed DH, Frankham R. 2003. Correlation between fitness and genetic diversity. Conservation Biology 17, 230–237. http://dx.doi.org/10.1046/j.1523-1739.2003.01236.x

Rodrigues L, Berg CVD, Povoa O, Monteiro A. 2013. Low genetic diversity and significant structuring in the endangered *Mentha cervina* populations and its implications for conservation. Biochemical Systematics and Ecology 50, 51–61. http://dx.doi.org/10.1016/j.bse.2013.03.007

Setoguchi H, Mitsui Y, Ikeda H, Nomura N, Tamura A. 2011. Genetic structure of the critically endangered plant *Tricyrtis ishiiana* (Convallariaceae) in relict populations of Japan. Conservation Genetics 12, 491–501. http://dx.doi.org/10.1007/s10592-010-0156-y

Slatkin M. 1994. Gene flow and population structure, In: Real LA (ed), Ecological Genetics, Princeton University Press, Princeton, NJ, USA, pp 3–17.

Stern FC. 1946. A study of the genus Paeonia. The Royal Horticultural Society, London.

Su X, Zhang H, Dong LN, Zhang JQ, Zhu XT, Sun K. 2006. RAPD classification and identification of *Paeonia rockii* varieties planted in Gansu Province. Acta Botanica Boreali-Occidentalia Sinica 26, 696–701.

Suo ZL, Zhang HJ, Zhang ZM, Chen FF, Chen FH. 2005. DNA molecular evidences of the inter-specific hybrids between *Paeonia rockii* and *P. suffruticosa* based on ISSR markers. Acta Botanica Yunnanica 27, 42–48.

Tan XJ, Wu ZK, Cheng WD. 2011. Association analysis and its application in plant genetic research. Chinese Bulletin of Botany 46, 108–118. http://dx.doi.org/10.3724/SP.J.1259.2011.00108

Tansley SA, Brown CR. 2000. RAPD variation in the rare and endangered *Leucadendron elimense* (Proteaceae): implications for their conservation. Biological Conservation 95, 39–48. http://dx.doi.org/10.1016/S0006-3207(00)00015-X

Tong F, Xie DF, Zeng XM, He XJ. 2016. Genetic diversity of *Paeonia decomposita* and *Paeonia decomposita* subsp *rotundiloba* detected by ISSR markers. Acta Botanica Boreali-Occidentalia Sinica 36, 1968–1976.

Varshney RK, Graner A, Sorrells ME. 2005. Genic microsatellite markers in plants: features and applications. Trends in Biotechnology 23, 48–55. http://dx.doi.org/10.1016/j.tibtech.2004.11.005

Wang DX, Ma H, Zhang YL, Duan AA, Li WJ, Li ZH. 2013. *Paeonia* (Paeoniaceae) expressed sequence tag-derived microsatellite markers transferred to *Paeonia delavayi*. Genetics and Molecular Research 12, 1278–1282. http://dx.doi.org/10.4238/2013.April.17.6

Wang JX, Xia T, Zhang JM, Zhou SL. 2009. Isolation and characterization of fourteen microsatellites from a tree peony (*Paeonia suffruticosa*). Conservation Genetics 10, 1029–1031. http://dx.doi.org/10.1007/s10592-008-9680-4

Wright S. 1978. Evolution and the genetics of populations. Vol. 4: Variability within and among natural populations, University of Chicago Press, Chicago.

Wu F, Zhang D, Ma JX, Luo K, Di HY, Liu ZP, Zhang JY, Wang YR. 2016. Analysis of genetic diversity and population structure in accessions of the genus *Melilotus*. Industrial Crops and Products 85, 84–92. http://dx.doi.org/10.1016/j.indcrop.2016.02.055

Xu XX, Cheng FY, Xian HL, Peng LP. 2016. Genetic diversity and population structure of endangered endemic *Paeonia jishanensis* in China and conservation implications. Biochemical Systematics and Ecology 66, 319–325. http://dx.doi.org/10.1016/j.bse.2016.05.003

Yang Y, Liu JK, Zeng XL, Wu Y, Song HX, Liu GL. 2015. A comparative study on composition of seed oil fatty acids of some wild populations of *Paeonia decomposita*. Acta Horticultural Sinica 42, 1807–1814.

Yang Y, Luo JT, Zhang BF, Song HX, Liu GL, Zeng XL. 2015. Studies on floral characteristics and breeding system of *Paeonia decomposita*. Journal of Plant Resources and Environment 24, 97–104.

Yao XH, Ye QG, Kang M, Huang HW. 2007. Microsatellites analysis reveals interpopulation differention and gene flow in endangered tree *Changiostyrax dolichocarpa* (Styracaceae) with fragmented distribution in central China. New Phytologist 176, 472–480. http://dx.doi.org/10.1111/j.1469-8137.2007.02175.x

Yu HP, Cheng FY, Zhong Y, Cai CF, Wu J, Cui HL. 2013. Development of simple sequence repeat (SSR) markers from *Paeonia ostii* to study the genetic relationships among tree peonies (Paeoniaceae). Scientia Horticulturae 164, 58–64. http://dx.doi.org/10.1016/j.scienta.2013.06.043

Yuan JH, Cheng FY, Zhou SL. 2012. Genetic structure of the tree peony (*Paeonia rockii*) and the Qinling Mountains as a geographic barrier driving the fragmentation of a large population. PLoS One 7, e34–55. http://dx.doi.org/10.1371/journal.pone.0034955

Zhang JJ, Shu QY, Liu ZA, Ren HX, Wang LS, Keyser, De E. 2012. Two EST-derived marker systems for cultivar identification in tree peony. Plant Cell Reports 31, 299–310. http://dx.doi.org/10.1007/s00299-011-1164-1

Zhang JM, López-Pujol J, Gong X, Wang HF, Vilatersana R, Zhou SL. 2018. Population genetic dynamics of Himalayan-Hengduan tree peonies, Paeonia subsect. Delavayanae. Molecular Phylogenetics and Evolution 125, 62–77. http://dx.doi.org/10.1016/j.ympev.2018.03.003

Zhang JS, Yang CY, Wu C, Hu ZY, Wang RG, Guo YS, Ren XL. 2012. Study on genetic diversity, population structure and specificity of subpopulations of fluecured tobacco germplasm. Acta Tabacaria Sinica 18, 21–49.

Zhang P, Zhou ZC, Jin GQ, Fan HH, Hu HB. 2006. Genetic diversity analysis and provenance zone allocation of *Schima superba* in China using RAPD markers. Scientia Silvae Sinicae 42, 38–42.

Zhang YL, Han XY, Niu LX, Zhang J, He LX. 2015. Analysis of fatty acid in seed oil from nine wild peony species. Journal of Chinese Cereals and Oils Association 30, 72–79.

Zhao X, Zhou ZQ, Lin QB, Pan KY, Li MY. 2008. Phylogenetic analysis of *Paeonia* sect. *Moutan* (Paeoniaceae) based on multiple DNA fragments and morphological data. Journal of Systematics and Evolution 46, 563–572. http://www.jse.ac.cn/EN/10.3724/SP.J.1002.2008.06197

Zhao X, Zhou ZQ, Lin QB, Pan KY, Hong DY. 2004. Molecular evidence for the interspecific relationships in *Paeonia* section *Moutan*: PCR-RFLP and sequence analysis of glycerol-3-phosphate acyltransferase (GPAT) gene. Acta Phytotaxonomica Sinica 42, 236–244.

Zhou ZQ, Pan KY, Hong DY. 2003. Advances in studies on relationships among wild tree peony species and the origin of cultivated tree peonies. Acta Horticuhurae Sinica 30, 751–757.

Zong XX, Guan JP, Gu JJ, Wang H, Ma Y. 2009. Differentiationon population structure and genetic diversity of pea core collections separately constituted from Chinese land races and international genetic resources. Journal of Plant Genetics and Resources 10, 347–353.

Zou YP, Cai ML, Wang ZP. 1999. Systematic studies on *Paeonia* sect. Moutan DC. based on RAPD analysis. Acta Phytotaxonomica Sinica 37, 220–227.

